# An investigation into the origins and history of pandemic small ruminant lentivirus infection

**DOI:** 10.1101/236117

**Authors:** Maria-Luisa Carrozza, Anna-Maria Niewiadomska, Maurizio Mazzei, Mounir R. Abi-Said, Stéphane Hué, Joshua B. Singer, Joseph Hughes, Robert J. Gifford

**Author notes:** Equal contributions (Maria-Luisa Carrozza, Anna-Maria Niewiadomska).

## Abstract

Small ruminant lentiviruses (SRLVs) cause chronic, persistent infections in populations of domestic sheep and goats throughout the world. In this study, we use genomic data to investigate the origins and history of the SRLV pandemic. To explore the hypothesis that SRLV infection disseminated during Neolithic times, we performed a serology and DNA sequencing-based investigation of SRLVs diversity in the Fertile Crescent region, where domestication of sheep and goats is thought to have originally occurred. While we found an elevated level of viral genetic diversity compared to other regions of the world, we did not find unambiguous evidence that the Fertile Crescent region was the centre of the contemporary SRLV pandemic. We therefore examined historical reports to investigate the relationship between contemporary SRLV distribution and diversity and the emergence of SRLV-associated disease. Historical data suggested that the emergence of SRLV-associated disease might be associated with the long-distance export of exotic small ruminant breeds - in particular, karakul sheep from Central Asia - during the late 19^th^ and early 20^th^ centuries. Phylogeographic analysis could neither confirm nor refute this hypothesis. However, we anticipate that future accumulation of genomic data from SRLV strains found throughout the world may allow for a more definitive assessment. The openly available data and resources assembled in this study will facilitate future investigations in this area.

**Importance:** Viruses that cause chronic, persistent infections have circulated in animals for millions of years. However, many have only emerged as pathogens within the far shorter timeframe of recorded human history. It is important to understand the history of chronic viral infections in domestic animals, so that more effective control and eradication programs can be developed.

## Introduction

Small ruminant lentiviruses (SRLVs) are retroviruses that infect domestic sheep (*Ovis aries*) and goats (*Capra hircus*), causing chronic, persistent infections that ultimately lead to organ failure and death. Infection is usually only apparent following an incubation period of ∼3-4 years, and lifelong subclinical infections have been reported (1). General susceptibility and specific disease manifestations are influenced by genetic factors, and vary between species and breeds, but typically include pneumonia, wasting, paralysis, polyarthritis and mastitis.

The prototypic SRLV isolate was obtained from an epidemic that emerged in Icelandic sheep in the late 1930s. This isolate was cultivated *in vitro* in the late 1950s, and was named ovine maedi-visna virus (OMVV) (1–4). A virus closely related to OMVV was isolated from a North American dairy goat in 1974 and named *caprine arthritis-encephalitis virus* (CAEV) (5). It has subsequently become clear that each of these viruses occurs in both sheep and goats. Accordingly, they are now considered to represent two distinct genotypes (A and B respectively) within of a virus species (SRLV) (6). Analysis of complete viral genomes indicates that at least five major SRLV genotypes circulate in domesticated small ruminants (A, B, B3, C and E) (7–9). Genotypes A and B are widespread throughout the world (6, 7, 10, 11) having apparently been disseminated by recent livestock trade (6). By contrast, the divergent genotypes B3, C and E appear to have relatively restricted distributions. Phylogenetic studies have implicated livestock trade in pandemic spread of SRLV genotypes A and B (6). However, the early events that gave rise to pandemic SRLV spread are not well understood. In this study, we examined SRLV genome data using molecular phylogenetic approaches to investigate the origins and early history of the SRLV pandemic.

## Materials and methods

### Serology and sequencing

Serological testing of sheep and goat flocks in Lebanon was conducted during August 2012. Sampling took place in various regions of the ‘Beqaa valley’, a local center of agricultural production. Flocks were chosen at random from the Central, West and Eastern Beqaa, as well as the Northern region of Mount Lebanon. The small ruminant population in Lebanon consists mainly of local fat-tailed ‘Awassi’ breed sheep, whereas the goat population is ∼95% ‘Baladi’ and ∼5% ‘Damascus’. Herds are often semi-nomadic, grazing during the day and returning to sheltered structures during the night, and frequently changing location depending on the season. Goats and sheep are often housed in the same areas and will graze together as one flock. Serological testing (ELISA) was conducted using plasma samples obtained from 449 animals: 221 goats and 228 sheep. A total of 36 goat, sheep and mixed flocks were sampled, with ten to fifteen animals in each flock being selected at random for plasma sampling. Of the initial 449 animals tested in Lebanon, ∼15.6 % (70 samples in 26 flocks: 20 goat; 50 sheep) were seropositive for exposure to SRLVs. Serotype was determined for a representative subset (n=45) of positive samples using P16-25 ELISA. Whole blood was spun at 3500rpm, the serum and buffy coat collected, and stored at −20C for further testing. ELISA assays were performed as described in (12). Net absorbance was obtained by subtracting the absorbance of negative antigen from the absorbance of each recombinant antigen. Cut-off value was defined as percentage of reactivity 20% of the absorbance of positive control included in each plate. Genomic DNA was extracted from stored white blood cells (buffy coats) using the DNeasy Blood and Tissue kit (QIAGEN), and quantitated using a nanodrop. Nested PCR was performed on the Gag region of the genome as described previously (13). PCR products were TA cloned into pCR4.0 sequencing vector (Life Technologies) and sent for sequence analysis (Genewiz). Viral sequences generated in this study have been submitted to GenBank under accession numbers KU170752-KU170766.

### Dataset construction

Previously published SRLV sequences were retrieved from GenBank in 2015 using the following search phrases to query the ‘Organism’ field: ‘Maedi Visna virus’; ‘Ovine Progressive Pneumonia Virus’; ‘Caprine Arthritis Encephalitis Virus’; ‘Small Ruminant Lentivirus’; ‘Ovine lentivirus’; ‘Caprine lentivirus’. We filtered the 5,255 sequences recovered via this means to derive a subset of 585 previously published sequences that each represented a distinct infection. For each determined the pubmed ID of the study it was obtained in, the isolate name, the year of isolation, and the host species (sheep or goat). This information was obtained from GenBank files where possible, or else from scientific papers associated with the published sequences, and failing that via direct contact with study authors. Sequences in this dataset were aligned using a combination of MUSCLE (14) and Pal2Nal (15). We used GLUE – a data-centric bioinformatics framework for working with virus sequence data (16) – to capture the relationships between sequences, alignments and associated tabular data utilised in our investigation. The resultant resource is publicly available via GitHub (https://github.com/giffordlabcvr/SRLV-GLUE).

### Literature search/historical investigation

To investigate previous reports of SRLV outbreaks, we conducted electronic searches for studies written in English, German, French, Portuguese, and that documented SRLV outbreaks based on either pathology, serology; by nucleic acid amplification. In November 2012, we searched the following databases: PubMed/Medline, JSTOR, Web of Science, WorldCat and Google scholar. In addition searches were conducted for outbreaks reported in ProMED Mail, as well as for sequences from unpublished studies reported in the National Center for Biotechnology Information’s nucleotide database. A non-redundant list of reports was obtained using the following search terms; ‘Maedi Visna Virus’; ‘Visna’; ‘Maedi’; ‘Ovine Progressive Pneumonia Virus’; ‘Caprine Arthritis Encephalitis Virus’; ‘MVV’; ‘CAEV’; ‘SRLV’; ‘Small Ruminant Lentiviruses’; ‘Montana Sheep Disease’; ‘Montana Progressive Pneumonia’; ‘Chronic Progressive Pneumonia’; ‘Zwoegerziekte’.

### Phylogenetic, phylogeographic and phylodynamic analysis

Phylogenetic analysis of SRLV *gag* sequences obtained from sheep in Lebanon and of complete SRLV genome sequences was performed using RaXML (17), using the general time reversible (GTR) nucleotide substitution model and 1000 bootstrap replicates. Time-calibrated phylogenies were reconstructed using BEAST 1.8.2 with the SDR06 substitution model and an uncorrelated lognormal relaxed clock model. Molecular clock model comparison was investigated using AICM available in Tracer. In all cases the model comparison indicated support for the relaxed clock. Additionally, the posterior distribution for the coefficient of variation in the uncorrelated lognormal relaxed clock model did not impinge on zero indicating that the relaxed clock model provides a better fit to the data than the strict clock. After testing for the best molecular clock model, we tested two different coalescent models: Bayesian skyline plot and Bayesian skyride. Trees were annotated using FigTree (http://tree.bio.ed.ac.uk).

The dispersal between countries was estimated using the discrete state phylogeographic approach described in Lemey et al. (18) using a symmetric substitution model and implementing the Bayesian stochastic search variable selection model (BSSVS). The Bayesian GMRF skyride coalescent model was selected as a flexible demographic tree prior. Stationarity was assessed on the basis of the effective sampling size (ESS) after a 10% burning using Tracer and two independent runs were generated to confirm convergence. The two independent runs were combined using LogCombiner and summarized with TreeAnnotator. In cases where there was a large amount of uncertainty in the node heights resulting in negative branch length, the common ancestor tree was used to summarise the trees instead of the Maximal Clade Credibility topology. Whole genome analysis did not produce ESS values above 100 for the root height and a number of other parameters for SRLV-A and SRLV-B probably due to insufficient data.

## Results

### SRLV diversity in the Fertile Crescent region

Archaeological and genetic evidence indicate that both sheep and goats were domesticated in the ‘Fertile Crescent’ region of Western Asia ∼10.5-11 thousand years ago (19) (Figure 1a). The presence of highly divergent SRLV genotypes in remnant populations of primitive small ruminant breeds (genotype C in Norway, genotype E in Sardinia) is consistent with an ancestral radiation of SRLV genotypes out of the Fertile Crescent region in association with the diffusion of early agricultural systems during the early Neolithic (20, 21). However, this pattern might also be accounted for in other ways (e.g. each genotype might represent an independent introduction of virus into the domesticated small ruminant population from a wild ruminant species).

**Figure 1.**
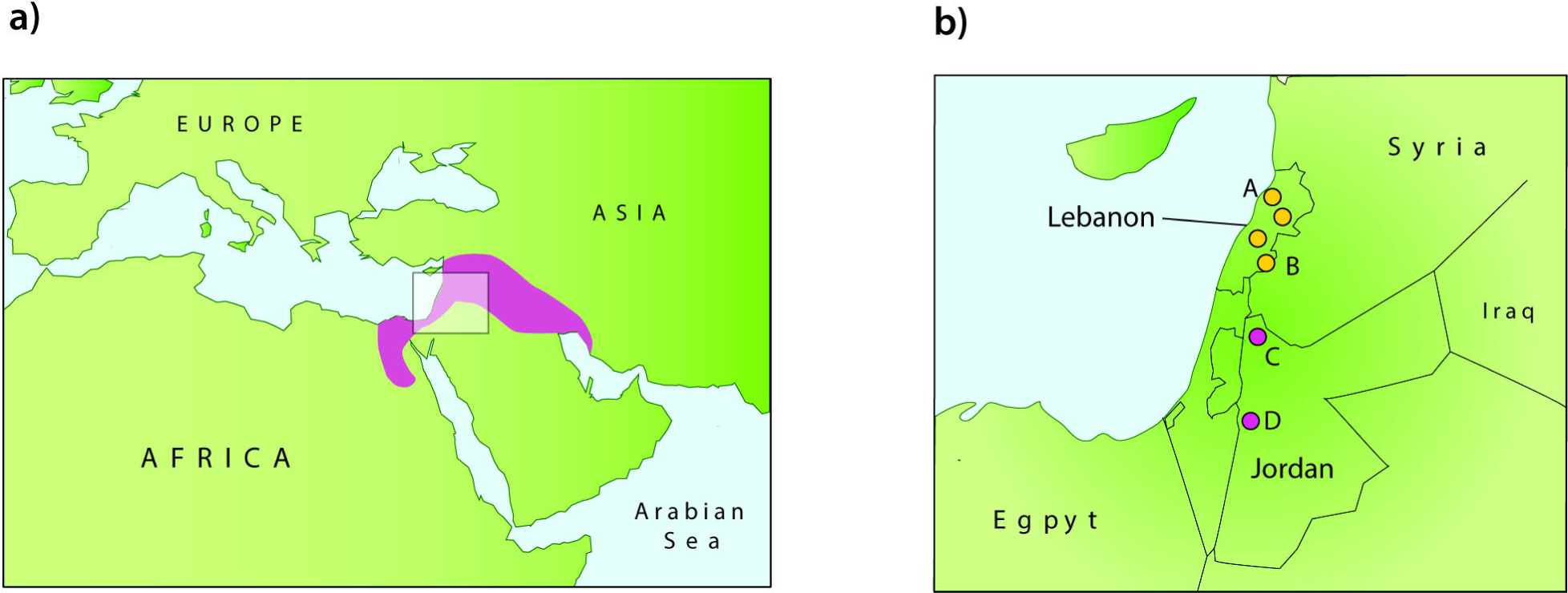
Sampling in the Fertile Crescent region. The map in panel **(a)** indicates the ‘Fertile Crescent’ region in pink. The overlapping square indicates the location of the area shown in panel **(b)**, which includes the countries sampled in this study. Yellow circles indicate sites in Lebanon where sampling was performed in this study; A=Northern Lebanon (Qornet el Sawda, Arez); B= Beqaa Valley. Beqaa Valley comprised three sites; North (Aammiq), West (Swairi, Manara, Rashaya); Eastern (Nahle, Maqne, Knaisse). Pink circles indicate sites previously sampled in Jordan: C=Northern Jordan; D=Jordan Valley.

We carried out a study of SRLV diversity in the Fertile Crescent region to investigate whether this area might have been the centre of an ancient SRLV radiation. We focused on Lebanon, a country in the Levant region, where (Figure 1b), where some of the earliest human civilizations are believed to have developed (22). Serological testing of 886 animals indicated that diverse SRLVs are present in both countries (Table 1). Furthermore, overall prevalence was relatively high (∼21%) in comparison with previous global surveys (23), but similar to estimates obtained in other countries in the region (13, 24). We used PCR to obtain *gag* gene amplicons from a randomly selected subset of our SRLV-positive samples (primers are listed in **Table S1**).

**Table 1.** ELISA results from small ruminants in Jordan and Lebanon

| Species (Breed) | Sample number | Positive samples |  | Serotype reactivity |  |  |
| --- | --- | --- | --- | --- | --- | --- |
|  |  | # | % | A | B | E |
| Sheep (Awassi) | 457 | 131 | **28.7** | 23 | 20 | 9 |
| Sheep (Australian) | 5 | 3 | **60.0** | 2 | - | 1 |
| Goat (Baladi) | 408 | 48 | **11.8** | 11 | 24 | 4 |
| Goat (Damascus) | 14 | 4 | **29.0** | 1 | 1 | 1 |
| Goat (Baladi x Damascus) | 2 | 0 | **0.00** | - | - | - |
| **Totals** | **886** | **186** | **21** | **37** | **45** | **15** |

**Legend:** SRLV prevalence and genotypes in Jordan and Lebanon. ELISA assays were performed on sera collected from sheep and goats belonging mostly to local breeds. The serotype reactivity of a subset of sera comprising 45 Lebanese and 112 Jordanian samples was analysed with a genotyping ELISA assay targeted against the p16/matrix (MA) and p25/capsid (CA) proteins of A, B and E genotype.

We obtained amplicons from fifteen samples and investigated their evolutionary relationships to a representative set of published sequences (Figure 2). In phylogenies constructed using this dataset, three distinct clades of Levantine isolates (Lev I-III) were observed. The Lev-I and Lev II clades grouped robustly within the diversity of previously characterized SRLV-A isolates, while Lev-III was relatively distinct from all previously sampled SRLV genotypes. A full-length proviral genome sequence has previously been recovered for one Lev-III isolate sampled in Jordan (Jord1). We used full-length genome sequences to reconstruct the phylogenetic relationships between Jord1 and other SRLV genomes, showing it is approximately as distantly related to established SRLV genotypes as they are to one another (**Figure S1**).

**Figure 2.**
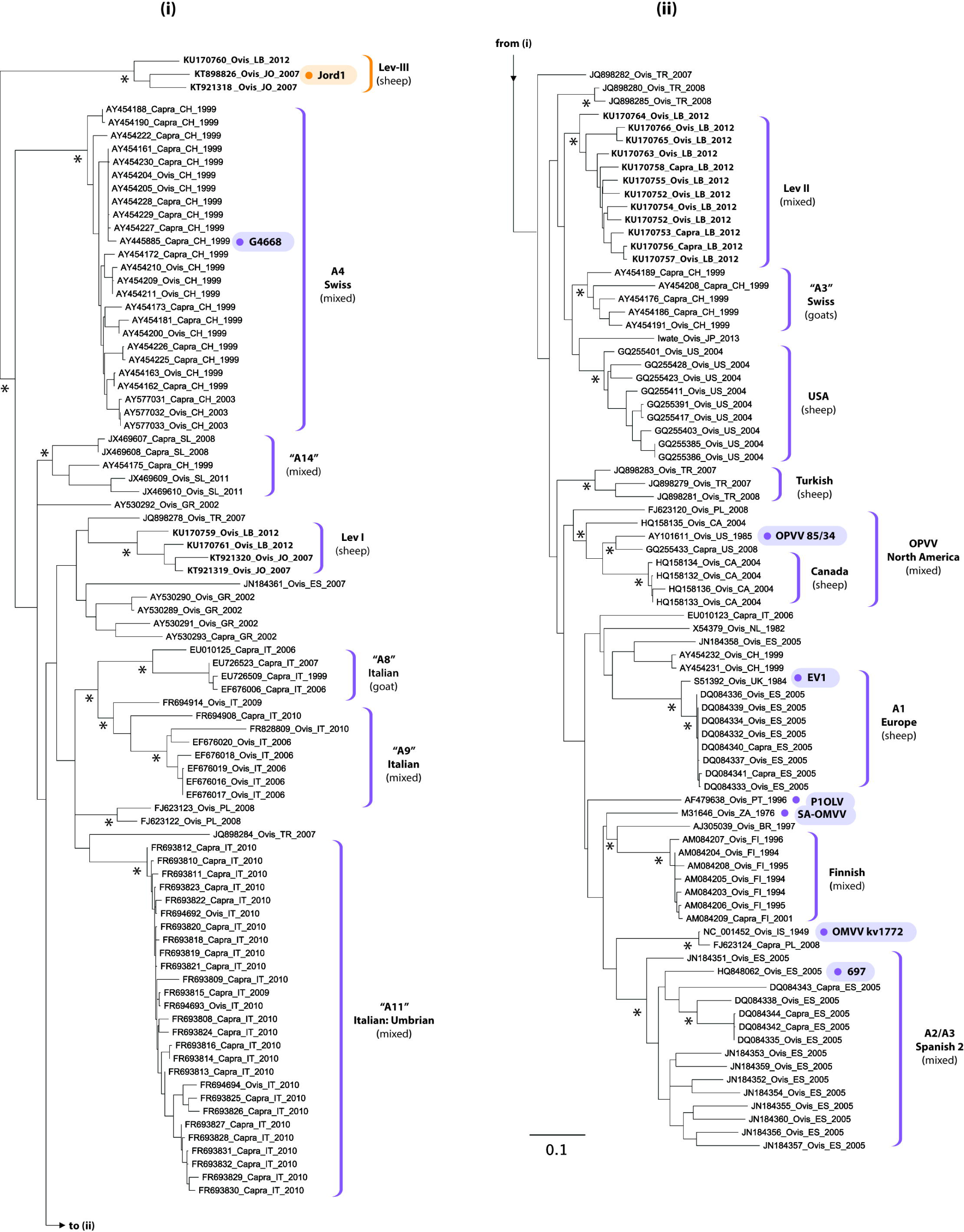
Maximum likelihood (ML) phylogeny of 219 SRLV sequences based on capsid (CA) gene nucleotide sequences. The tree is midpoint rooted, and has been split across two panels. Taxa labels include isolate details separated by underscores as follows; accession number/isolate; species (Ovis=*Ovis aries*; Capra=*Capra hircus*); country of sampling (ISO two-letter country code); year of sampling. Circles next to taxa names indicate complete genome sequences, strain is shown in bold for these isolates. Brackets on the right indicate clades with bootstrap support that either derive from a single country or geographic region, or that have previously been assigned genotype subgroup status. For these clades, text associated with each bracket indicated whether the clade has been found in sheep only, goats only, or both sheep and goats (mixed). Inverted commas indicate that subgroup status is based on analysis of sub-genomic sequences. Asterisks indicate nodes with ML bootstrap support >85%, based on 100 bootstrap replicates. The scale bar indicates evolutionary distance in substitutions per site.

While the Lev III clade is highly divergent, it does not represent an obvious ancestor to all other SRLV genotypes. Furthermore, although the overall level of diversity in the Fertile Crescent region appears to be relatively high, based on observations from our study and others (10), the range of genotypes and novel lineages identified was not as exceptionally high as might be expected if this region were the centre of an ancient SRLV radiation (i.e. similar to the high diversity of human immunodeficiency virus type 1 (HIV-1) found in areas of West Central Africa). In fact, most of the isolates detected in our study grouped within the diversity of pandemic SRLV-A strains found in Europe and North America, indicating they are as likely to have been imported into the region as exported from it (Figure 2).

### Recorded history of pandemic SRLV emergence and spread

Previous epidemiological studies have indicated that pandemic spread of SRLVs may only have occurred relatively recently (6, 23, 25, 26). They do not appear to have spread in parallel with the first European colonists, as seems to be the case for many other livestock pathogens (27, 28). For example, SRLV-A has never been reported in Australia or New Zealand, where sheep are largely descended from animals imported in the 19th century from the UK, Germany, South Africa and North America (6, 25). This suggests that SRLVs were not widespread during the ‘Age of Discovery’ (from ∼1500 to ∼1700), when the first exports of small ruminants from Europe to Africa, Oceania and the Americas took place.

To explore the manner in which SRLVs emerged as globally distributed pathogens, we reviewed the historical evidence for SRLV-associated disease in countries throughout the world. For each country where SRLV infection has been reported, we determined the earliest reliable evidence for the presence of the virus (**Table S1, Table S2**). Previous studies have identified diseases reported in the past that appear likely - in retrospect - to have been caused by SRLVs (1). Cases where the only available evidence for historical SRLV infection is based on a description of disease pathology have to be viewed cautiously, particularly because of potential confusion with sheep pulmonary adenomatosis, caused by Jaagsiekte sheep retrovirus (JSRV). Nonetheless, SRLV-A can confidently be stated to have been present in Europe since at least 1933, because the Icelandic epidemic from which the prototypic SRLV-A strain was derived was definitively traced to an importation of Karakul sheep in this year (29).

Intriguingly, the early emergence of SRLV-A infection closely mirrors the early 20^th^ century diffusion of Karakul, which are among the oldest domesticated sheep breeds (30, 31) (Figure 3). The breed is native to Central Asia, and arrived in Iceland via the Institute for Animal Breeding, in Halle, Germany. Karakul sheep were originally exported to Halle from Bukhara in Uzbekistan in 1903 (32), and were also introduced to the United States in the same decade (33) (**Table S1**). In the first half of the 20^th^ century, a cluster of similar disease syndromes emerged, all of which are now considered likely to have been caused by SRLV infection (29). ‘*Graaf-Reinet disease’* was first described in South Africa in 1915, less than a decade after the introduction of Karakul into the region. The same year, ‘*Montana sheep disease’* was reported for the first time in the United States, seven years after the arrival of Karakul. In Europe, both *maedi-visna* (in Iceland), and ‘*la bouhite’* (in France) emerged in ∼1930-50, in parallel with the initial dissemination of German Karakul in Europe.

**Figure 3.**
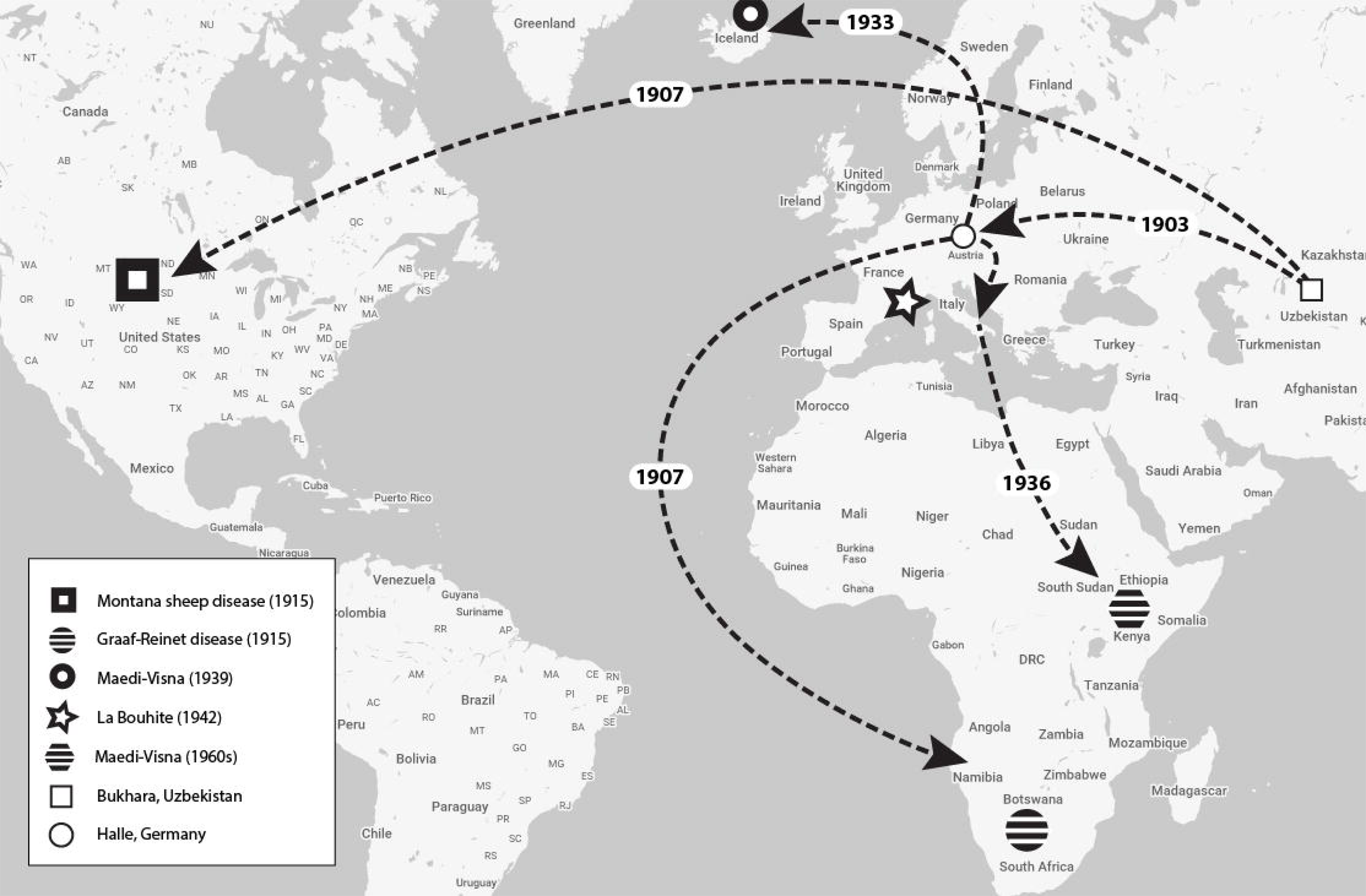
Map showing the geographic spread of Karakul in relation to early outbreaks of infection. Dashed lines show export of Karakul sheep in the period 1900-1950. Early outbreaks of disease associated with SRLV infection are indicated (see key), as recorded in **Table S2**.

Similarly, we observed that the pandemic spread SRLV-B (specifically subtype B1) was conspicuously associated with Swiss dairy goat breeds, particularly the Swiss Saanen breed (**Table S2**). Many of the first reported outbreaks of SRLV infection in goats involved this popular breed (34). Indeed, the prototypic SRLV-B isolate (CAEV Cork) derives from an outbreak of chronic, progressive disease that emerged in a North American population of Saanen goats in 1970 (35). Until the late 19^th^ century, small ruminant breeds such as Saanen and Karakul had strictly defined geographic distributions. A role for the livestock trade in SRLV emergence has been proposed previously. However, the role of specific breed movements in driving the early emergence of pandemic strains has not been closely examined.

### Phylogeographic investigation of pandemic SRLV spread

We used Bayesian phylogeographic approaches to investigate whether available genomic data supported a role for breed dispersal in the emergence of pandemic SRLV infection, as suggested by our investigation of historical records. We compiled a comprehensive set of published SRLV sequences. From an initial set of 5,255 sequences, we derived a subset of 697 that together represented 585 unique field isolates, each obtained from a distinct infected animal (Table 2, **Table S3**). We added the fifteen SRLV *gag* sequences we obtained in Lebanon to this set. Together, this set of sequences represented 600 isolates sampled in a total of 30 countries, with dates of sampling ranging from 1949 to 2013. A preliminary phylogenetic analysis (data not shown) was performed to determine the genotype of all sequences in this set. We then created individual multiple sequence alignments (MSA) for each of the pandemic SRLV strains (SRLV-A and SRLV-B1). From these alignments, we derived a total of eight alignment partitions (**Table S4**) - two within the group-specific antigen (*gag*) gene, and two partitions within the polymerase (*pol*) gene. We detected temporal structure in five of the eight datasets, despite the presence of outliers (**Table S5**). The results of Bayesian phylogeographic analysis of the four *pol* partitions are shown in Figure 4 (SRLV-A) and Figure 5 for (SRLV-B1), while the results obtained for the four *gag* partitions are shown in **Figure S2** (SRLV-A) and **Figure S3** for (SRLV-B1). Table 4 and **Table 5** provide information about specific nodes of interest in all eight phylogenies. We obtained more robust results when using *pol*, however, we have included the results obtained for *gag* as these provide some potentially useful insights regarding the epidemic in specific parts of the world.

**Table 2.** Summary of SRLV sequence studies

| Study <sup>a</sup> | Accessions (strain) <sup>b</sup> | Species <sup>c</sup> | Date <sup>d</sup> | Genomic region(s) <sup>e</sup> | Country <sup>f</sup> | # Seqs <sup>e, g</sup> |  |
| --- | --- | --- | --- | --- | --- | --- | --- |
|  |  |  |  |  |  | Set | Study |
| S. America |  |  |  |  |  |  |  |
| (43) | AJ305039-42 | Ch, Oa | 1989-97 | Gag | BR | 4 | 4 |
| DS | AF402664-66, 68 | Ch | 1999 | Gag | BR | 4 | 4 |
| DS | AY101347-8 | Ch | NK | Gag | BR | 2 | 2 |
| DS | AY081139 | Ch | NK | Gag | BR | 1 | 1 |
|  |  |  |  |  |  | 11 |  |
| N. America |  |  |  |  |  |  |  |
| (44) | HQ158122-36 | Ch, Oa | 2004 | Gag | CA | 15 | 15 |
| (45) | HM210570 (FESC-752) | Ch | 2004 | Genome | MX | 1 | 1 |
| (45) | HM212792-3 | Ch | 2008 | Env | MX | 2 | 2 |
| (46) | K03327 | Ch | 1985 | Pol | US | 1 | 1 |
| (47, 48) | NC_001463 (Cork) | Ch | 1974 | Genome | US | 1 | 1 |
| (49) | See legend. | Oa | 2004, 08 | Gag | US | 10 | 52 |
| (50) | See legend | Oa | 1976, 03 | Env | US | 8 | 17 |
| (5, 51) | M60855 | Ch | 1980 | Env | US | 1 | 1 |
| (52), DS | AY101611 (85/34) | Oa | 1985 | Gag-Pol | US | 1 | 1 |
| (52, 53) | U64439 (85/34) | Oa | 1985 | Env | US | 1 | 1 |
|  |  |  |  |  |  | 41 |  |
| Africa |  |  |  |  |  |  |  |
| (54) | M31646 (SA-OMVV) | Oa | 1976 | Genome | ZA | 1 | 1 |
| DS | FJ619565-72 | Ch | 2007 | Gag | SU | 8 | 8 |
|  |  |  |  |  |  | 9 |  |
| East Asia |  |  |  |  |  |  |  |
| (55) | GU120138 (Shanxi) | Ch | 1995 | Genome | CN | 1 | 1 |
| (56) | AY900630 (Gansu) | Ch | 1992 | Genome | CN | 1 | 1 |
| (41) | AB821356 | Oa | 2013 | Gag | JP | 1 | 1 |
| (40) | N/A | Ch | N/K | Gag | KO | 1 | 1 |
| DS | AB747557, LC002526 | Ch | 2011 | Gag | PH | 1 | 2 |
| DS | GU903321 | Ch | 2009 | Env | TW | 1 | 1 |
| DS | HM237197 | Ch | 2010 | Env | TW | 1 | 1 |
| DS | See legend | Ch | 2008-11 | Gag | TH | 14 | 14 |
|  |  |  |  |  |  | 20 |  |
| West Asia |  |  |  |  |  |  |  |
| (57) | JQ898278-85 | Oa | 2007-08 | Gag | TR | 8 | 8 |
| (10) | JF502418 (TR-DM) | Oa | 2006 | Gag | TR | 1 | 1 |
|  |  |  |  |  |  | 9 |  |
| South West Europe |  |  |  |  |  |  |  |
| (58)* | See supp. | Oa | 2008-11 | Env | ES | 8 | 350 |
| (59) | JN184351-78 | Oa | 2008-11 | Gag | ES | 28 | 28 |
| (59) | JN184379-94 | Oa | 2008-11 | Env | ES | 16 | 16 |
| (60) | DQ084332-44 | Ch, Oa | 2005-06 | Gag | ES | 13 | 13 |
| (60) | DQ632733-35 | Ch, Oa | 2005-06 | Pol | ES | 3 | 3 |
| (61) | FJ195346 (Ov496) | Oa | 2004 | Genome | ES | 1 | 1 |
| (62) | HQ848062 (697) | Oa | 2005 | Genome | ES | 1 | 1 |
| (63) | AF479638 (P1OLV) | Oa | 1996 | Genome | PT | 1 | 1 |
|  |  |  |  |  |  | 71 |  |
| North West Europe |  |  |  |  |  |  |  |
| (64) | AM084203-09 | Oa | 1994-96, 01 | Gag | FI | 7 | 7 |
| (64) | AM084210-15 | Ch, Oa | 1994-96, 01 | Env | FI | 6 | 6 |
| (65) | U35796, 798-801, 804, 811 | Ch, Oa | 1994 | Env | FR | 7 | 7 |
| (66) | DQ149845 | Oa | 2004 | Env | FR | 1 | 1 |
| (67) | AJ400718-21 | Ch | 1994 | Env | FR | 4 | 4 |
| (68) | AF015181 | Ch | 1994 | Gag | FR | 1 | 1 |
| (68) | AF015182 | Ch | 1994 | Pol | FR | 1 | 1 |
| (68) | AF015180 | Ch | 1994 | Env | FR | 1 | 1 |
| DS | AJ969032-37 | Ch | 1998-02 | Pol | IE | 6 | 6 |
| DS | AJ969038-43 | Ch | 1998-02 | Env | IE | 6 | 6 |
| (69) | U51910 | Oa | 1962 | Env | IS | 1 | 1 |
| (70, 71) | NC_001452 (kv1772) | Oa | 1949 | Genome | IS | 1 | 1 |
| DS | X54379 | Oa | 1970 | Gag | NL | 1 | 1 |
| (72) | DQ844911-30 | Ch, Oa | 1991-03 | Pol | NO | 20 | 20 |
| (72) | DQ844931-41 | Oa | 1976-05 | Env | NO | 11 | 11 |
| (20) | DQ015910-16 | Ch | 1991-02 | Env | NO | 7 | 7 |
| (20) | AF322109 (1GA) | Ch | 1991 | Genome | NO | 1 | 1 |
| (73) | S51392 (EV1) | Oa | 1984 | Genome | UK | 1 | 1 |
|  |  |  |  |  |  | 84 |  |
| Central, South East and East Europe |  |  |  |  |  |  |  |
| (6) | AY454161-232 | Ch, Oa | 1999 | Gag | CH | 72 | 72 |
| (6) | AY454233-296 | Ch, Oa | 1999 | Pol | CH | 64 | 64 |
| (6) | AY445885 (**G4668**) | Ch | 1999 | Genome | CH | 1 | 1 |
| (7) | AY577031-3 | Ch, Oa | 2003 | Gag-pol | CH | 3 | 3 |
| (7) | AY577034-9 | Ch, Oa | 2003 | Gag | CH | 6 | 6 |
| (74) | AY530289-93 | Ch, Oa | 2002 | Gag | GR | 5 | 5 |
| (75)* | HM449450 | Ch | 2007 | Env | IT | 1 | 234 |
| (76)* | EF685749, 83 | Ch | 2006 | Env | IT | 2 | 194 |
| (11) | *See legend* | Ch, Oa | 2009-10 | Gag-pol | IT | 55 | 61 |
| (77)* | EF144472, 82, 93, 94 | Ch | 2006 | Env | IT | 4 | 45 |
| (37) | EU293537 (**Roccaverano**) | Ch | 2008 | Genome | IT | 1 | 1 |
| (37) | EU726488-525 | Ch | 1999, 06, 07 | Gag | IT | 38 | 38 |
| (8) | EF676000-26 | Ch, Oa | 2006 | Gag | IT | 24 | 24 |
| (8) | EU010120-6 | Ch | 2006 | Gag | IT | 7 | 7 |
| (78) | DQ013214-43 | Ch, Oa | 2005 | Pol | IT | 30 | 30 |
| (79) | AY044803-10 | Ch | 1991 | Gag | IT | 8 | 8 |
| (80) | GQ381130 (**Seui**) | Ch | 2003 | Genome | IT | 1 | 1 |
| (80) | GQ428519-36 | Ch | 2008-09 | Gag | IT | 18 | 18 |
| (10) | JF502416 (**Fonni**) | Oa | 2003 | Genome | IT | 1 | 1 |
| (10) | JF502417 (**Volterra**) | Oa | 2004 | Genome | IT | 1 | 1 |
| (10) | JF520393-406 | Ch, Oa | 2008-09 | Gag | IT | 5 | 5 |
| (81) | EU709743, EU702487 | Oa | 1991 | Env | IT | 2 | 3 |
| (82) | FJ623120-25 | Ch, Oa | 2008 | Gag | PL | 6 | 6 |
| (82) | FJ623110-19 | Ch, Oa | 2008 | Env | PL | 10 | 10 |
| DS | JN084103 | Ch | 2008 | Gag | RU | 1 | 1 |
| DS | JN008914 | Ch | 2008 | Env | RU | 1 | 1 |
| (83) | JQ610956-JQ611030 | Oa | 2011 | Pol | SL | 75 | 75 |
| (83) | JX469600-10 | Ch, Oa | 2008-11 | Gag-pol | SL | 11 | 11 |
|  |  |  |  |  |  | **453** |  |
| Total |  |  |  |  |  | **697** |  |

**Legend.** Previously sequenced SRLV isolates, grouped by study. <sup>a</sup> DS = direct submission to GenBank. Asterisks indicate studies of intra-host diversity from which a subset of sequences (one per animal) was selected. <sup>b</sup> Accession numbers are shown only for sequences included in the reference set. For complete genome sequences the isolate name is shown in bold and in parenthesis. NA = not applicable. <sup>c</sup> Oa=Ovis aires; Ch=Capra hircus. <sup>d</sup> Year of collection is shown where it could be ascertained, NK = not known. <sup>e</sup> Only sequences that encoded >500 base pairs of gag, pol,; env were included in the reference set. <sup>f</sup> ISO country code (state). <sup>g</sup> Reference set was restricted to one sequence per infected animal (number shown as 'in set'). Where multiple sequences were available the total is indicated under 'in study'. Accession numbers not listed in the table are as follows; Thai sequences (GQ161209-15, JF714253-5, EU983108-9, EU919141, FJ167525)); North American sequences from reference (49) (GQ255391, GQ255386, GQ255401, GQ255411, GQ255385, GQ255403, GQ255417, GQ255423, GQ255428, GQ255433); North American sequences from reference (50) (AY362022, AY362024, AY362026, AY362029, AY362030, AY362032, AY362035, AY362038); Italian sequences from reference (11) (FR687200, FR693808-32, FR694686-96, FR694908-21, FR695719, FR828809-16).

**Table 4.** tMRCAs of selected clades within SRLV-B

| Partition | Node # | Clade description | Posterior | Region(s) | Location | Loc P | Age | 95% HPD |
| --- | --- | --- | --- | --- | --- | --- | --- | --- |
| Gag MSA 1 | 1 | Thailand-Poland | 1.0 | C. Europe / S.E. Asia | Thailand | 0.77 | 1993 | 10-33 |
| Gag MSA 1 | 2 | Spanish | 1.0 | W. Europe | Spain | 0.97 | 1992 | 13-29 |
| Gag MSA 1 | 3 | Italian | 0.84 | W. Europe | Italy | 0.94 | 1990 | 11-36 |
| Gag MSA 1 | 4 | TH-PL-ES-IT | 0.75 | Europe / S.E. Asia | Italy | 0.71 | 1981 | 20-45 |
| Gag MSA 1 | 5 | Canada-China-Mexico | 0.62 | N. America | Canada | 0.71 | 1982 | 21-42 |
| Gag MSA 1 | 6 | Global | 0.84 | Global | Canada | 0.37 | 1978 | 25-42 |
| Gag MSA 1 | 7 | Root | 1.0 | Global | USA | 0.73 | 1972 | 37-54 |
| Gag MSA 2 | 1 | Slovenia-Italy | 0.83 | S. Europe | Switzerland | 0.79 | 1888 | 51-206 |
| Gag MSA 2 | 2 | Brazil | 1.0 | S. America | Brazil | 1.0 | 1974 | 19-60 |
| Gag MSA 2 | 3 | Canada-Mexico | 0.92 | N. America | Canada | 1.0 | 1877 | 61-228 |
| Gag MSA 2 | 4 | USA-Canada-Mexico-CN | 0.85 | Global | Canada | 0.5 | 1856 | 70-263 |
| Gag MSA 2 | 5 | Switzerland-Italy | 0.83 | Europe | Switzerland | 1.0 | 1862 | 66-250 |
| Gag MSA 2 | 6 | Root | 1.0 | Global | Switzerland | 0.92 | 1825 | 82-322 |
| Pol MSA 1 | 1 | Italy-Switzerland | 1.0 | C. Europe | Switzerland | 0.98 | 1986 | 16-34 |
| Pol MSA 1 | 2 | Slovenia | 1.0 | S. Europe | Slovenia | 1.0 | 2001 | 6-12 |
| Pol MSA 1 | 3 | Italy-Switzerland | 0.96 | Europe | Switzerland | 1.0 | 1966 | 25-47 |
| Pol MSA 1 | 4 | Root | 1.0 | Global | Switzerland | 0.63 | 1973 | 37-53 |
| Pol MSA 2 | 1 | Slovenia | 1.0 | S. Europe | Slovenia | 1.0 | 1989 | 11-35 |
| Pol MSA 2 | 2 | Switzerland-Slovenia | 0.89 | Europe | Switzerland | 1.0 | 1963 | 35-61 |
| Pol MSA 2 | 3 | China-US-Switzerland | 0.89 | Global | Switzerland | 1.0 | 1960 | 44-59 |
| Pol MSA 2 | 4 | Root | 1.0 | C. Europe | Switzerland | 1.0 | 1952 | 47-72 |

**Figure 4.**
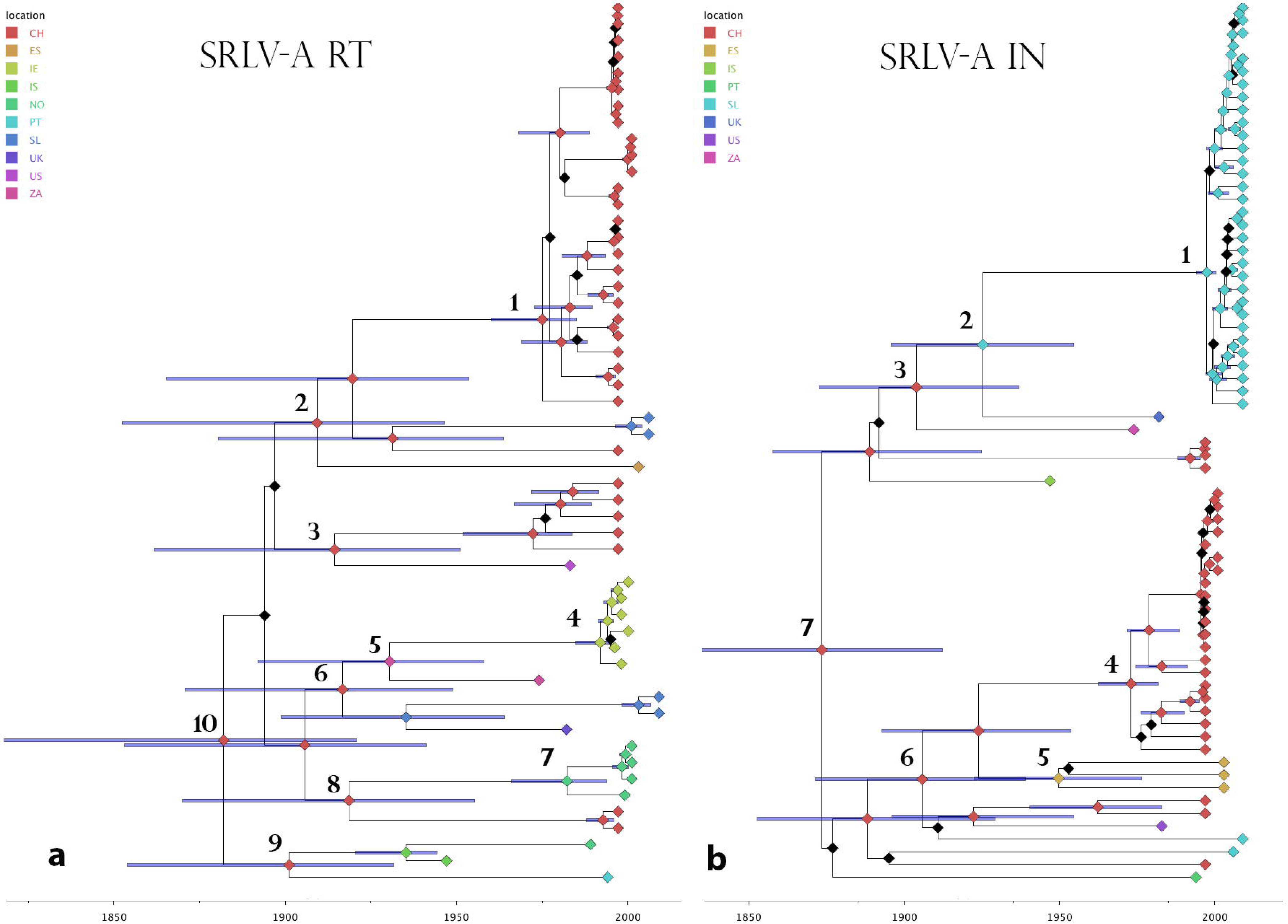
Time-calibrated maximum likelihood Bayesian phylogenies of SRLV-A. Panel **(a)** shows analysis of an MSA partition in reverse transcriptase (RT), panel **(b)** shows analysis of an MSA partition in integrase (IN). The scale bar indicates year dates. The key to the upper left shows colour codes for the predicted country location of each node. Nodes for which a country estimate could not be derived with posterior probabilities <0.5 are coloured black. Error bars illustrating mean coalescence time estimates with 95% highest posterior density (HPD) are shown on internal nodes.

**Figure 5.**
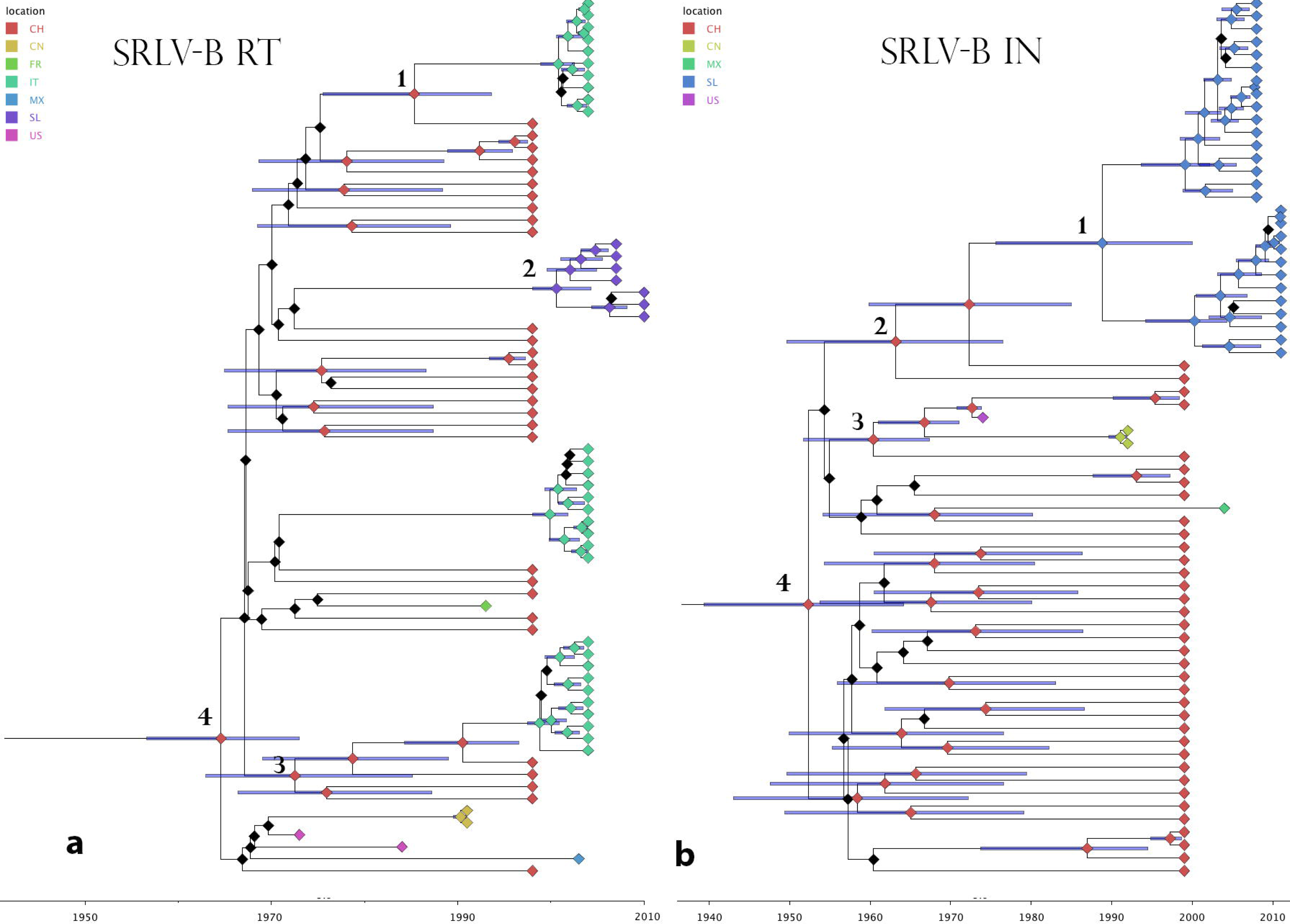
Time-calibrated maximum likelihood Bayesian phylogenies of SRLV-B1. Panel **(a)** shows analysis of an MSA partition in reverse transcriptase (RT), panel **(b)** shows analysis of an MSA partition in integrase (IN). The scale bar indicates year dates. The key to the upper left shows colour codes for the predicted country location of each node. Nodes for which a country estimate could not be derived with posterior probabilities <0.5 are coloured black. Error bars illustrating mean coalescence time estimates with 95% highest posterior density (HPD) are shown on internal nodes.

In general, phylogenies were only moderately structured as opposed to ‘star-like’. We obtained robust support for nodes near the phylogeny tips, particularly within clades of viruses sampled from the same geographic region, but only a proportion of the deeper branching relationships (i.e. between clades of viruses sampled in distinct counries and/or geographic regions) had high support. For all four *pol* partitions, and one SRLV-B1 *gag* partition, the location of the most recent common ancestor was placed in Switzerland. Ancestral locations within Asia Minor were obtained for SRLV-A *gag* partitions, but with low support. One SRLV-B1 gag partition placed the most recent common ancestor in the United States (US), with moderately high support.

Overall, the results of Bayesian analysis were consistent with a scenario within which both SRLV-A and SRLV-B1 disseminated worldwide from a European epicentre. Where we were able to confidently estimate the location of deeper nodes in the phylogeny, we found these were placed in Central Europe. By contrast, clades sampled in Asia, South America and the around the Northern and Western periphery of Europe (i.e. Ireland, Norway, Finland) were observed to have relatively recent common ancestors (Table 4, **Table 5**).

Although the lack of resolution in our phylogenies (as well as the limitations of opportunistic sampling) prevented us from inferring the precisely the direction and timing of global SRLV spread, we were able to confidently estimate tMRCAs for most of the MSAs examined. For SRLV-A, we obtained tMRCA estimates in the late 19^th^ century - shortly predating the first Karakul exports from Central Asia - for both reverse transcriptase (RT) (1883, 1820-1923 95%HPD) and integrase (IN) (1887, 1837-1915 95%HPD). The SRLV-A *gag* partitions yielded more conflicting results: for the matrix partition (Gag MSA 1) we obtained much older estimates with wide error margins, whereas the capsid partition (Gag MSA 2) yielded a somewhat more recent tMRCA (Table 3). For SRLV-B1 the capsid partition yielded unreliable tMRCA estimates with wide error margins. However, we obtained more reliable and generally consistent results from the other three MSA partitions, which provided tMRCA estimates of 1966 for RT (1958-1974 95%HPD), 1952 for IN (1939-1964 95%HPD), and 1972 for matrix (1957-1974 95 HPD) (Table 4).

**Table 3.**
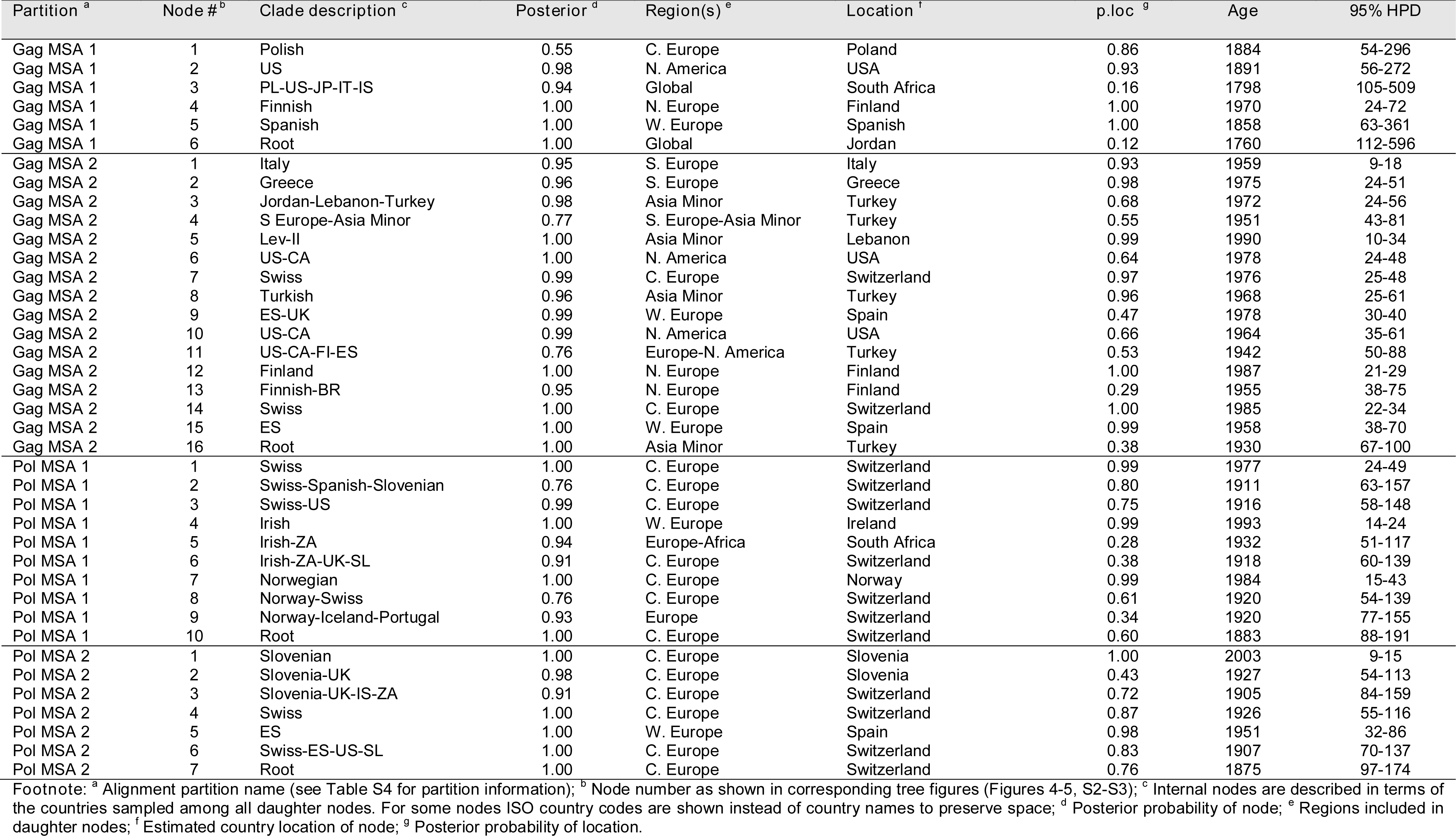
tMRCAs of selected clades within SRLV-A

## Discussion

In this study, we used molecular phylogenetic approaches to investigate the origins and history of pandemic SRLV spread. To investigate the hypothesis that SRLV originally disseminated out of Western Asia during the early Neolithic period (i.e. in association with domesticated small ruminants) we sampled SRLV strains circulating in the Fertile Crescent region where domestication of these species is thought to have occurred. We found a relatively high prevalence and diversity of SRLV strains, but we did not find obvious evidence of the region having been the centre of an ancient SRLV radiation. The lack of clear evidence that the various SRLV genotypes derive directly from a virus population that circulated in the ancestral small ruminant populations from which modern breeds are derived, does not rule out the possibility that they did. It could be that our sampling was not entirely representative, or that SRLV diversity in the region has been greatly influenced in recent times by factors such transport and migration (6).

To investigate the origins and timing of pandemic SRLV spread, we combined a review of historical records with a phylogenetic investigation of SRLV genome data. Our research led us to consider the possibility that the emergence of pandemic SRLV infection might have its roots in the late 19^th^ and early 20^th^ century, during which time ‘exotic’ small ruminant breeds were exported to Europe, North America and Africa. In particular, we noted that historical reports suggest the early pattern of emergence of SRLV-associated disease around the world appears – at least superficially – to be correlated with the export of karakul – a breed of fat-tailed sheep, native to Central Asia (Figure 3). Even though it was established relatively early on that SRLV-A infection was imported to Iceland along with a flock of karakul sheep in the 1930s, this breed has not previously been implicated in global spread of SRLV. However, a role for karakul in the early emergence of SRLV-A might easily have been missed if the SRLV strains exported along with this breed were not associated with obvious disease in their natural host. Current evidence indicates that Lentiviruses, including some SRLVs, are relatively apathogenic in populations with which they have longer-term associations (36, 37). Thus, in the light of what we now know lentivirus pathology and its relationship to host genetic variation, a plausible hypothesis accounting for the emergence of pandemic SRLV-associated disease is that it was driven by the introduction (via karakul export) of Central Asian SRLV strains into populations of small ruminants in Europe, Africa and America that had previously only been exposed to SRLV strains prevalent in Europe.

We propose that the early spread of SRLVs could have been enabled by the creation of new contact networks in livestock populations, leading to particular small ruminant breeds and populations being brought into contact for the first time. Subsequent to this, the development of international livestock trade (6) - as well as the development and uptake of new systems of agriculture - would presumably have helped facilitate the further spread of SRLVs from European and Asian epicenters to Africa, the Americas, Oceania, and East Asia. The capacity of some small ruminant breeds to harbor SRLV infection without exhibiting symptoms, and a lack of understanding about the nature of slow virus infections prior to the 1950s (29), would presumably helped enable this relatively recent wave of global SRLV spread.

To investigate whether available genomic data supported this hypothesis, we performed a phylogeographic analysis of the available sequence data for the pandemic SRLV strains A and B1. Overall, results were consistent with both strains having spread worldwide relatively recently (i.e. within the 20^th^ century), emerging from a Eurasian source. For SRLV-A, we obtained tMRCA estimates in the late 19^th^ century - shortly predating the first Karakul exports from Central Asia – and thus consistent with a scenario under which these exports ultimately led to the emergence of an SRLV pandemic.

SRLV-B1 appears to have disseminated globally more recently than SRLV-A, from an unknown source in Europe. Global spread of this SRLV lineage is conspicuously associated with Swiss dairy goat breeds in general, and the Saanen breed in particular (**Table S2**). Many of the first reported outbreaks of SRLV infection in goats involved this breed, which was first exported from Switzerland in the late 19^th^ century, and has subsequently been introduced to many countries throughout the world (34). However, given that (i) SRLV infection is known to cause disease in Saanen and other Swiss dairy goat breeds, and; (ii) these breeds have been exported from Switzerland since the late 19^th^ century (34), It seems unlikely that Swiss dairy goats are the ancestral hosts of SRLV-B1. One possibility is that spread of B1 from a Swiss epicenter was preceded by the introduction of novel SRLV diversity into the Swiss small ruminant population during the early 20^th^ century.

A major limitation of our analysis is that the majority of sequence data we examined was opportunistically sampled from what was already available in GenBank. Sampling of SRLV diversity so far has been patchy, and in addition, most of the available sequences are sub-genomic (see Table 2), which limits what can be inferred. Furthermore, sampling is heavily biased toward certain areas, distorting our phylogeographic inference. For example, Switzerland was probably the most thoroughly sampled country in our dataset, thus the phylogeographic placement of the SRLV-A and SRLV-B MRCAs in Switzerland has to be viewed with some caution. Interestingly, however, Karakul are documented as having been present in Switzerland since at least the early 1930s (38).

Test and removal programs are currently used as a means of controlling SRLV infection, with varying levels of effectiveness (39). In fact, recent years have seen the emergence of SRLV infection in countries where it had not previously been documented, as well as the re-emergence of disease in areas where it had been considered to be under control (40, 41). We therefore anticipate that the use of genome data to investigate SRLV breakdowns will increase in future. The increased availability of genome sequences from SRLV strains found throughout the world – particularly in Central Asia – should eventually allow for a more definitive assessment of the hypothesis presented here, and provide a clear picture of the pattern of pandemic SRLV spread. The more ancient spread of SRLV viruses, which is proposed to date back to the Neolithic, is unlikely to be amenable to phylogeographic analysis due to the time-dependence of viral evolutionary rates (42). However, the overall distribution of SRLV strains in Europe and Asia (particularly Central Asia) may well be revealing in this respect. The resources assembled in this study will not only facilitate further investigation of historical SRLV spread, but can also provide a foundation for the development of SRLV surveillance systems that utilise genomic data.

## Acknowledgements

RJG was supported by a grant from the UK Medical Research Council (No. MC_UU_12014/10). We thank Elie Barbour for contributing materials, lab space and equipment at the University of Beirut, Francesco Tolari for providing Jordanian sheep and goat blood samples, and Roman Biek, Pablo Murcia, Massimo Palmarini and Tiago Saraiva for useful discussions. We also thank Valgerður Andrésdóttir, Katerina Angelopoulou, Beatriz Amorena, Barbara Blacklaws, Miguel Fevereiro, Gordon Harkiss, Isidro Hötzel, Jinhai Huang, Britt Gjerset, Donald P. Knowles, Caroline Leroux, Bandit Nuansirchay, Keisuke Oguma, Monica Olech, Ramsés Reina, Sergio Rosati, Jörg Schüpbach, Hiroshi Sentsui, Hyun-Jin Shin, and Stephen Valas for providing SRLV sequence information.

